# SignalP 6.0 achieves signal peptide prediction across all types using protein language models

**DOI:** 10.1101/2021.06.09.447770

**Authors:** Felix Teufel, José Juan Almagro Armenteros, Alexander Rosenberg Johansen, Magnús Halldór Gíslason, Silas Irby Pihl, Konstantinos D. Tsirigos, Ole Winther, Søren Brunak, Gunnar von Heijne, Henrik Nielsen

## Abstract

Signal peptides (SPs) are short amino acid sequences that control protein secretion and translocation in all living organisms. As experimental characterization of SPs is costly, prediction algorithms are applied to predict them from sequence data. However, existing methods are unable to detect all known types of SPs. We introduce SignalP 6.0, the first model capable of detecting all five SP types. Additionally, the model accurately identifies the positions of regions within SPs, revealing the defining biochemical properties that underlie the function of SPs *in vivo*. Results show that SignalP 6.0 has improved prediction performance, and is the first model to be applicable to metagenomic data.

SignalP 6.0 is available at https://services.healthtech.dtu.dk/service.php?SignalP-6.0

## Main Text

Signal peptides (SPs) are short N-terminal amino acid (AA) sequences that target proteins to the secretory pathway in eukaryotes and for translocation across the plasma (inner) membrane in prokaryotes. As experimental identification of SPs is costly, SP prediction is a well-established task with high relevance in biological research (1). SP prediction tools enable identification of proteins that follow the general secretory (Sec) or the twin-arginine translocation (Tat) pathway and predict the position in the sequence where a signal peptidase (SPase) cleaves the SP (2, 3). The current state-of-the-art algorithm, SignalP 5.0, is able to predict Sec substrates cleaved by SPase I (Sec/SPI) or SPase II (Sec/SPII, prokaryotic lipoproteins), and Tat substrates cleaved by SPase I (Tat/SPI) (4). However, it is unable to detect Tat substrates cleaved by SPase II or Sec substrates processed by SPase III (prepilin peptidase, sometimes referred to as SPase IV (2)). Such type Sec/SPIII SPs control the translocation of type IV pilin-like proteins, playing a key role in adhesion, motility and DNA uptake in prokaryotes (5). Furthermore, SignalP 5.0 is agnostic regarding the SP structure, as it cannot define the three subregions (n-region, h-region and c-region) that underlie the biological function of SPs.

Here we present SignalP 6.0, based on powerful protein language models (LMs) (6–8) that leverage information from millions of unannotated protein sequences across all domains of life. LMs create semantic representations of proteins which capture their biological properties and structure. Using these protein representations, SignalP 6.0 can predict additional types of SPs that previous versions have been unable to detect, while extrapolating better to proteins distantly related to those used to create the model and to metagenomic data of unknown origin. Additionally, SignalP 6.0 is capable of identifying the three subregions of SPs.

We compiled, to our knowledge, the most comprehensive dataset of protein sequences that are known to contain SPs. In total, we gathered 3,352 Sec/SPI, 2,261 Sec/SPII, 113 Sec/SPIII, 595 Tat/SPI, 36 Tat/SPII, 16,421 intracellular and 2,615 transmembrane sequences. Moreover, we defined region-labeling rules according to known properties of the respective SP type (Fig. 1A). We applied three-fold cross-validation to train and evaluate the model. In our data partitioning procedure, we ensured that homologous sequences are placed in the same partition to be able to accurately measure the model’s performance on unseen sequences (Fig. S2).

**Fig. 1.**
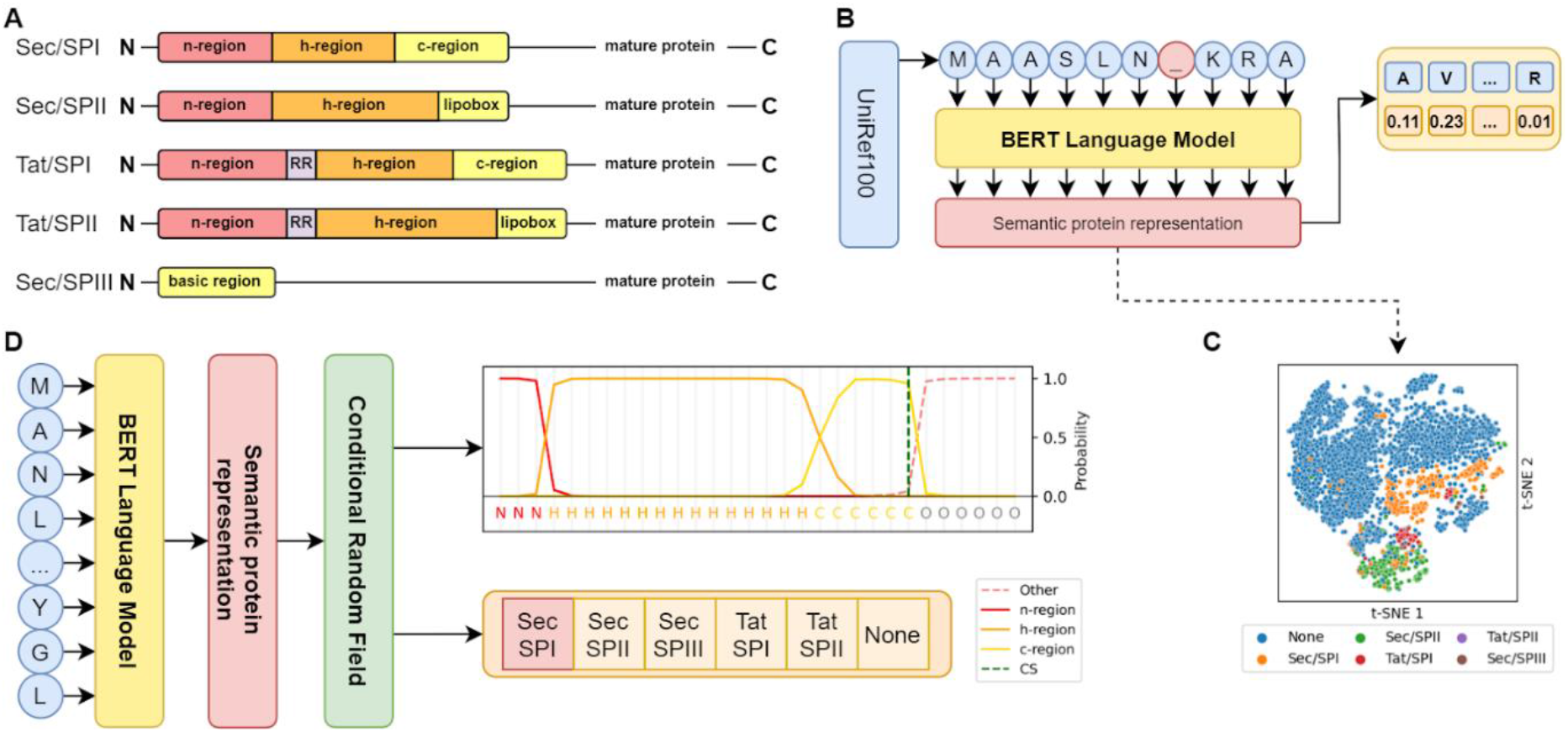
Modeling signal peptide structure using protein language models. **(A)** Region structures of the five types of SPs. Tat-translocated SPs feature a twin-arginine (RR) motif, SPs cleaved by SPase II a C-terminal lipobox. Sec/SPIII SPs have no substructure. **(B)** Protein LM training procedure. BERT learns protein features by predicting masked AAs in sequences from UniRef100. **(C)** t-SNE projection of protein representations before prediction training. Sequences with different SP types form distinct clusters, separated from sequences without SPs. **(D)** SignalP 6.0 architecture. An AA sequence is passed through the LM and the resulting semantic representation serves as input for the CRF. The CRF predicts the probability of each region at each position in conjunction with the SP type.

For previous predictors, Sec/SPIII and Tat/SPII were omitted due to a lack of annotated samples that makes it challenging for models to learn the defining features of these two types (4). Notably, this lack does not correspond to prevalence in nature, as these types exist throughout most organisms present in the databases (9, 10). Additionally, the available annotated sequences do not cover the full diversity encountered in nature, as they are biased towards well-studied organisms. Furthermore, existing predictors require data where the organism of origin is known, as this allows them to explicitly account for known differences in SP structure between Eukarya, Archaea, and Gram-positive and Gram-negative bacteria.

Recently, protein LMs have been shown to improve performance on problems with limited annotated data (11). Moreover, LM protein representations directly capture the evolutionary context of a sequence (6, 7). We hypothesized that using an LM, we would (I) obtain better performance on SP types with limited data availability, (II) achieve better generalization to sequences that are distantly related to training sequences and (III) enable prediction of sequences where the species of origin is not known. We opted for the BERT protein LM available in ProtTrans (6) that was trained on UniRef100 (12) (Fig. 1B). The LM was subsequently optimized on our dataset to predict SPs. We found that, already before optimization, the LM captured the presence of SPs in its protein representations (Fig. 1C). We combined the LM with a Conditional Random Field (CRF) probabilistic model (13) to predict the SP region at each sequence position in conjunction with the SP type, yielding the SignalP 6.0 architecture (Fig. 1D).

We evaluated SignalP 6.0 with nested cross-validation. As the baseline, we retrained the state-of-the-art architecture SignalP 5.0 on our new dataset. For all types except for Tat/SPI in Archaea, SignalP 6.0 shows improved performance. Especially for the two underrepresented types, Sec/SPIII and Tat/SPII, detection performance improves drastically (Fig. 2A), whereas performance of SignalP 5.0 remains too low to make it practically useful. This confirms the importance of LMs for low-data problems, making SignalP 6.0 the first model capable of simultaneously detecting all five types of SPs. Additionally, we find significant precision gains for predicting cleavage sites (Fig. 2B).

**Fig. 2.**
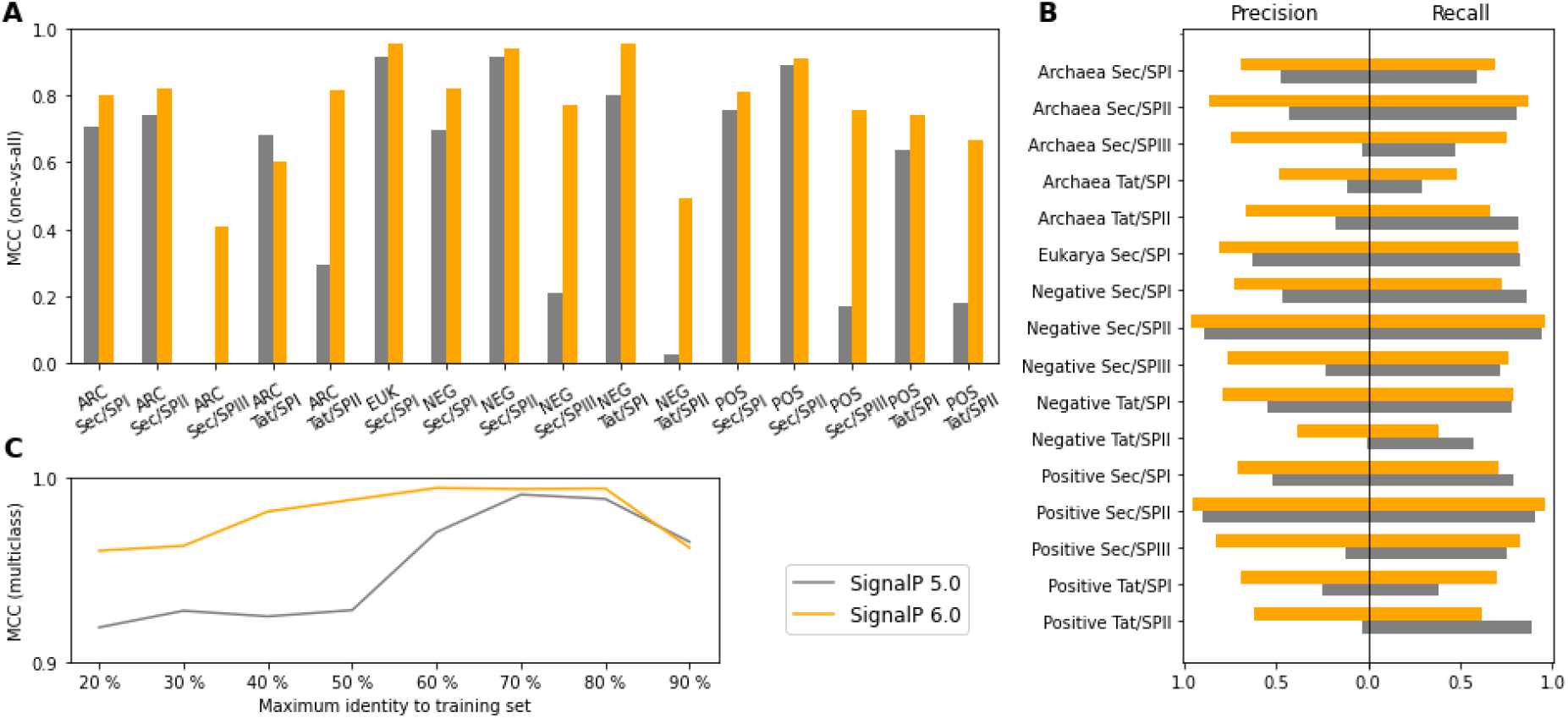
SignalP 6.0 shows strong performance on all types and organism groups. **(A)** SP detection performance measured as one-vs-all Matthews Correlation Coefficient. ARC=Archaea, EUK=Eukarya, NEG=Gram-negative bacteria, POS=Gram-positive bacteria. The positive set consists of the type, the negative set of non-SP sequences and all other types. SignalP 6.0 significantly improves performance on underrepresented types. **(B)** CS prediction performance. SignalP 6.0 has improved precision for all types and organism groups. **(C)** Dependence of performance on identity to sequences in the training data. At sequence identities lower than 60%, SignalP 6.0 outperforms SignalP 5.0.

We further benchmarked SignalP 6.0 against other publicly available predictors. In some cases, specialized predictors show stronger performance on the specific tasks they were optimized for (Tables S3-S8). However, none of these predictors are capable of detecting all SP types, and the results are further biased as they cannot be evaluated in a cross-validated setup.

When predicting a set of test sequences grouped by identity to any sequence in the training data, we find that detection performance at high sequence identities remains comparable. However, at identities lower than 60%, SignalP 6.0 outperforms SignalP 5.0, showing better generalization to proteins distantly related to the ones present in the training data (Fig. 2C).

To gain insight into the diversity of SP usage throughout evolution, we predicted all reference proteomes available in Uniprot (14) (Tables S9-h). Predictions confirm exceptionally high Tat/SPII frequencies in Halobacteria, even though the training dataset only contains 3 such sequences. Moreover, we also identified bacterial species with high Tat/SPII and Sec/SPIII frequencies. Among all species present in the data, the only organisms without predicted SPs are bacterial endosymbionts, indicating that protein translocation and export are indispensable to free-living organisms.

Most SP predictors require knowledge of a sequence’s organism group of origin for optimal performance (4, 15, 16). SignalP 6.0 does not suffer from a performance reduction when removing this information, indicating that the evolutionary context, as encoded in the LM representation, already captures the organism group (Fig. S3). Ultimately, this makes SignalP 6.0 the first multi-class SP prediction tool that is applicable to sequences of unknown origin, as is typically the case in metagenomic research. For context, 1.7% of UniProt release 2021_02, equaling 3.5 million sequences, have no organism specified.

Signal peptides are traditionally described as consisting of three regions (17, 18). As there is no experimental technique known that identifies region borders, there is no labeled data available for measuring performance. Thus, we benchmark our region identification by comparing the properties of predicted regions to known properties from literature, finding that the predictions of SignalP 6.0 match all expected properties (18). For n-regions, the model correctly recovers the average length and the differences between organism groups (17) (Fig. 3A). Predicted h-regions are less hydrophobic in Tat-translocated SPs than in Sec SPs, a property that is known to contribute to the selectivity of the pathways (19). While the c-region is generally uncharged, in Tat SPs it can contain positively charged residues to avoid recognition by the Sec system (20). SignalP 6.0 accurately captures this property, with the majority of Tat/SPI c-regions having a net charge of 0 or +1. The model also predicts negatively charged Tat/SPI c-regions, hinting at negative charges also possibly being suitable to hinder recognition.

**Fig. 3.**
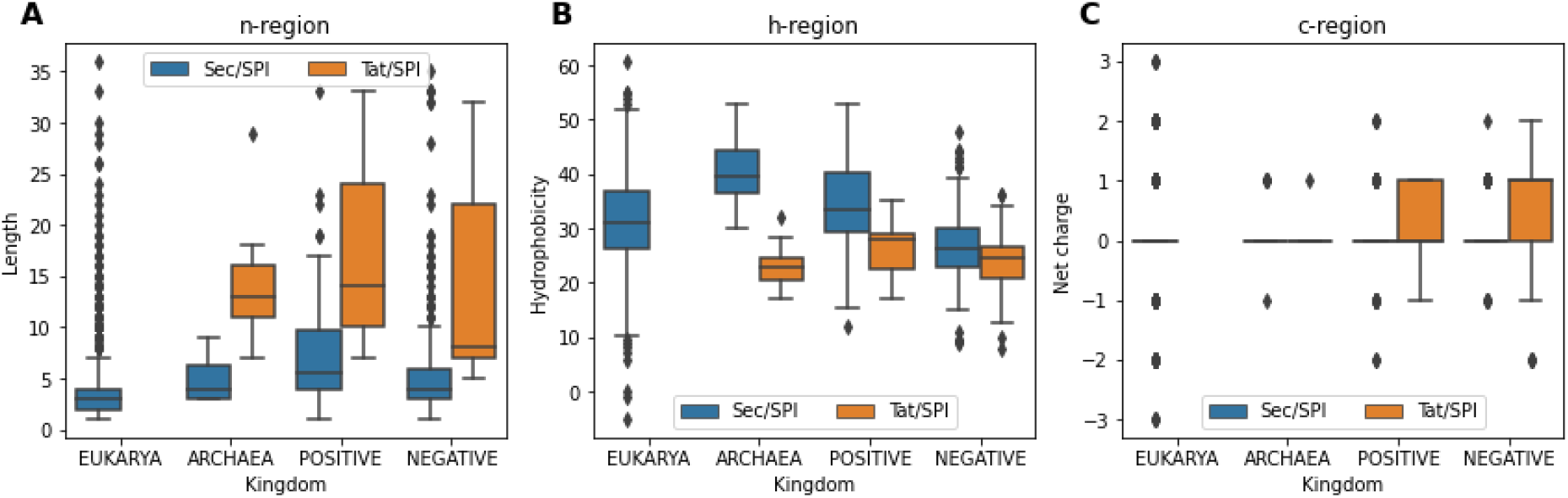
Predicted regions recapitulate known properties. **(A)** n-region lengths. The expected average length of about 4 residues for Sec/SPI SPs is recovered, n-regions are correctly predicted to be the shortest in eukarya and the longest in Gram-positive bacteria. **(B)** Hydrophobicities of the h-regions of Sec/SPI and Tat/SPI SPs. Sec-translocated SPs are predicted to have a higher hydrophobicity. **(C)** Net charges of Sec/SPI and Tat/SPI c-regions. Sec c-regions are uncharged, Tat c-regions are charged.

To further evaluate the region prediction capability, we predict a library of synthetic SPs that were found to be either functional or non-functional in *Bacillus subtilis* (21). In the original work, the authors did not find any discriminating properties between the two groups using traditional sequence analysis. Region predictions show a significant difference in n-region net charge (P<1×10^−4^) and hydrophobicity (P<1×10^−3^) between the groups (Fig. S4), revealing possible factors that contribute to in vivo functionality.

This study presents SignalP 6.0, the first model to achieve complete SP prediction by covering all five known types of SPs, while accurately predicting both sequences of unknown origin and evolutionarily distant proteins. Leveraging protein LMs, SignalP 6.0 is able to predict SP types with very limited training data available. By making the full spectrum of SPs accessible, the model allows us to further improve our understanding of protein translocation throughout evolution. In addition, identification of SP regions opens up new avenues into researching the defining properties that govern SP functionality. Given the potential of SPs as drug targets (22), and their emerging role in synthetic biology (21), investigating SPs and their properties at scale may lead to further advances in these fields.

## Materials and Methods

### Sequence data

The dataset for SignalP 6.0 was obtained by extending the data published with SignalP 5.0 (4). For all classes that were already part of the original data (Sec/SPI, Sec/SPII, Tat/SPI, soluble and transmembrane proteins), we added sequences that had become available in the respective source databases (UniProt (14) and Prosite (23) for signal peptides (SPs), UniProt and TOPDB (24) for soluble and transmembrane proteins) from 2018 until 7 November 2020, following the original selection criteria.

Tat/SPII sequences were identified using the combination of Prosite profiles PS51318 (Tat motif) and PS51257 (lipoprotein motif). By default, PS51318 is subject to post-processing that prevents both profiles from matching the same sequence. As there is experimental evidence for the existence of Tat-translocated lipoproteins (9, 10), we considered this post-processing rule to be biologically implausible. We disabled it manually in ScanProsite (25) and scanned all prokaryotic sequences in Swiss-Prot, yielding a total of 25 sequences where both profiles matched. Additional Tat/SPII sequences were found by training a simplified SignalP 6.0 model to discriminate SPII from non-SP sequences. We used this model to predict all Tat/SPI sequences in the training data, as we assumed that PS51257 is not sensitive enough to find all lipoproteins. We investigated the resulting hits in UniProt for supporting evidence of the proteins truly being lipoproteins, yielding 12 sequences that we relabeled to Tat/SPII from Tat/SPI. One additional sequence with manual evidence was found in the TatLipo 1.03 training data (9). For Sec/SPIII sequences we used Prosite pattern PS00409 for bacteria and Pfam (26) family PF04021 for Archaea, yielding 103 and 10 sequences respectively.

We improved the organism type classification of sequences by defining Gram-negative and Gram-positive bacteria more stringently, as we found that for edge cases such as Thermotogae, in which both gram stains can be observed (27), the classification in SignalP 5.0 was unclear. We redefined Gram-positive as all bacterial phyla that have a single membrane (monoderm): Actinobacteria, Firmicutes, Tenericutes, Thermotogae, Chloroflexi and Saccharibacteria. All remaining phyla have a double membrane (diderm) and were classified as Gram-negative.

We followed the methodology introduced by Gíslason et al. (28) for homology partitioning of the dataset into three partitions at 30% sequence identity. In brief, it achieves partitioning by computing the pairwise global sequence identities of all sequences using the Needleman-Wunsch algorithm (29), followed by single-linkage clustering. The resulting clusters are grouped together into the desired number of partitions. If there are sequences in a partition that have sequence identity higher than the defined threshold to a sequence in another partition, these sequences are iteratively removed until the maximum sequence identity criterion is fulfilled. The procedure was performed separately for each SP class, balancing for organism groups in each partition. The resulting partitions for each class were concatenated to yield the full dataset partitions.

The CD-HIT clustering method (30) that was employed in SignalP 5.0 enforces the homology threshold for cluster centers. However, as the training set was not homology reduced, but rather homology clustered, other data points can have a homology overlap significantly above the chosen threshold of 20% (Figure S2). When using Gíslason’s partitioning method, which strictly enforces the defined threshold, 20% maximum identity was impossible to achieve. Even at the relaxed threshold of 30%, the procedure resulted in the removal of a significant part of the dataset to achieve separation in three partitions (Table S3).

For benchmarking, we reused the benchmark set of SignalP 5.0, from which we excluded all sequences that were removed in the homology partitioning procedure of the new dataset. For sequences that were reclassified (to Gram-positive or to Tat/SPII), we changed the label accordingly.

For the synthetic signal peptide dataset, we used the data published by Wu et al. (21). We gathered all synthetic SP-mature protein pairs that were experimentally characterized, yielding 57 non-functional and 52 functional sequences. For the region analysis, we only considered sequences predicted as Sec/SPI signal peptides by SignalP 6.0, reducing the number of non-functional sequences to 55.

Reference proteomes and proteins of unknown origin were obtained from UniProt release 2021_02. To identify sequences of unknown origin, we used taxonomy identifiers 48479 (environmental samples), 49928 (unclassified bacteria) and 2787823 (unclassified entries).

### Generation of signal peptide region labels

We defined the task of learning signal peptide regions as a multi-label classification problem at each sequence position. Multi-label differs from multi-class in the sense that more than one label can be true at a given position. This approach was motivated by the fact that there is no strict definition of region borders that is commonly agreed upon, making it impossible to establish ground truth region labels for models to train on. We thus used the multi-label framework as a method for training with weak supervision, allowing us to use overlapping region labels during the learning phase that could be generated from the sequence data using rules. For inference, we do not make use of the multi-label framework, as we only predict the single most probable label at each position using Viterbi decoding, yielding a single unambiguous solution.

We defined a set of three rules based on known properties of the n-, h-, and c-regions. The initial n-region must have a minimum length of 2, and the terminal c-region a minimum length of 3 residues. The most hydrophobic position, which is identified by sliding a 7-AA window across the SP and computing the hydrophobicity using the Kyte-Doolittle scale (31), belongs to the h-region. All positions in-between these 6 labeled positions are labeled as either both n and h or n and c, yielding multi-tag labels.

This procedure was adapted for different SP classes, with only Sec/SPI completely following it. For Tat SPs, the n-h border was identified using the twin-arginine motif. All positions before the motif were labeled n, followed by 2 dedicated labels for the motif, again followed by a single position labeled as n. For SPII SPs, we did not label a c-region, as the C-terminal positions cannot be considered as such (17). The last 3 positions were labeled as the lipobox, all positions before that as h only. For SPIII SPs, no region labels were generated within the signal peptide.

### Modeling

SignalP 6.0 uses a pretrained protein language model (LM) to encode the amino acid sequence and a conditional random field (CRF) (13) decoder to predict the regions, cleavage sites and the sequence class labels. Specifically, we used the 30-layer Bert LM (32) that is available in ProtTrans (6), which was pretrained on UniRef100 (12). We removed the last layer of the pretrained model and extended the pretrained embedding layer by 4 additional randomly initialized vectors to represent the tokens for the 4 organism group identifiers. We prepend the organism group identifier to each sequence *s* of length *T* and encode it. From the resulting sequence of hidden states, we trim the positions corresponding to the organism group token and the special tokens used by BERT (CLS, SEP) to obtain a sequence of hidden states *h* of equal length as the original AA input *x*.

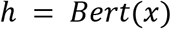

The hidden states serve as input for a linear-chain CRF. The CRF models the conditional probability of a sequence of states *y* = *y*_1_… *y_t_* given a sequence of hidden states *h* = *h*_1_… *h_t_* using the following factorization:

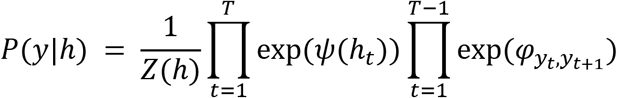

*Z*(*h*) is the normalization constant of the modeled distribution. *φ* is the learnable transition matrix of the CRF with *C* × *C* parameters, with *C* being the number of states (labels) modeled by the CRF. *ψ* is a learnable linear transformation that maps from the dimension of the hidden state h to the number of CRF states *C*, yielding the emissions for the CRF.

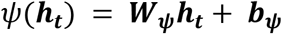

For each class of signal peptide G, there are multiple possible CRF states, corresponding to the defined regions of the SP class. We constrained the transitions in *φ* to ensure that regions are predicted in the correct order, leading to the possible state sequences depicted in Figure S1.

For inference we compute both the most probable state sequence (using Viterbi decoding) and the marginal probabilities at all sequence positions (using the forward-backward algorithm). The most probable state sequence is used to predict the cleavage site, which is inferred from the last predicted SP state as indicated in Figure S1.

As each SP consists of multiple regions, multiple states of C belong to a single global sequence class G. To predict the global class probabilities, we sum the marginal probabilities of all states that belong to a given class and divide the sum by the sequence length. This transforms a matrix of probabilities of shape C × T to a G × 1 vector of global class probabilities.

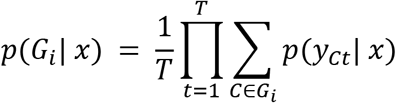

### Training

For training we minimize the negative log likelihood (NLL) of the CRF. As we can have multiple true labels *y_t_* at a given position, we use an extension of the equation known as multi-tag CRF. Multiple labels are handled by summing over the set of true labels *M_t_* at each position.

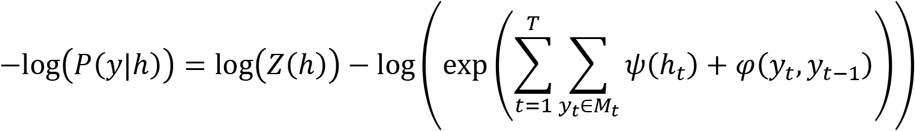

As we designed our region labels to be overlapping, the model is free to distribute its probability mass in any ratio between the correct labels at a given position. There are thus multiple solutions for the specific borders of n-, h-, and c- regions that yield the same NLL but are not equally biologically plausible. For instance, the model could learn a solution where it uniformly predicts an n-region of length 2 in all SPs, irrespective of the actual sequence. We employ regularization to promote the finding of biologically plausible solutions. Our regularization is based on the fact that the three SP regions have divergent amino acid compositions, which we can quantify by computing the cosine similarity between the amino acid distributions.

The most obvious approach would be to compute the amino acid distribution of each region based on the region borders inferred from the predicted most probable path of the sequence. This however cannot be used for regularization, as we require the term to be differentiable, which our Viterbi decoding implementation is not. We therefore based our regularization term on the marginal probabilities of the CRF computed by the forward-backward algorithm, which are used to compute a score for each amino acid for each region, approximating the discrete AA distributions.

For each region *r* ∈ {*n, h, c*}, we sum the marginal probabilities of all CRF states *c* belonging to region *r* at position *t*, yielding *S_t,r_*. We sum *S_t,r_* of all positions *t* of the sequence that have amino acid *a*, yielding the elements of the score vector ***score_r_*** for each region. We compute the cosine similarity between the normalized score vectors of *n* and *h* and *h* and *c*.

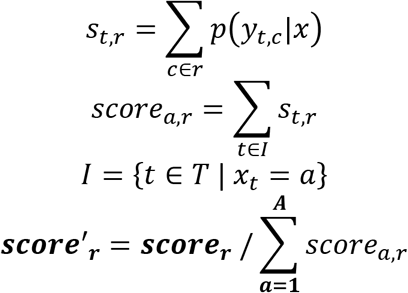

We perform this operation for each sequence. Sequences where a region does not exist (e.g. no c-region in Sec/SPII) are ignored for the respective similarity. The mean over all sequences for both similarities, multiplied by a factor *α*, was added to the loss. We observed that for about half the random seeds we tested, training runs with regularization enabled converged to a n-region length of 2 after one epoch. This is a degenerate solution, as this causes the n-region AA distribution to only be nonzero at a single position, yielding low similarity scores while being biologically implausible (a length of 2 is expected as the minimum, not the average over all sequences). Such runs were stopped and discarded after one epoch.

The model was trained end-to-end including all layers of Bert for 15 epochs, using Adamax as the optimizer and a slanted triangular learning rate. We applied dropout on the hidden state outputs of Bert to avoid overfitting. Hyperparameters were optimized using SigOpt (33). We employed three-fold nested cross-validation (outer loop is three-fold and inner loop is two-fold), yielding a total of 3 × 2 models for evaluation.

### Evaluation and benchmarking

For comparability, we employed the same metrics that were used in SignalP 5.0. SP detection performance was measured using the Matthews correlation coefficient (MCC) (34). We computed the MCC twice, once with the negative set only consisting of transmembrane and soluble proteins (MCC1), and once with it additionally including sequences of all other SP types (MCC2). For cleavage site (CS) prediction, we computed the precision and the recall. The precision was defined as the fraction of correct CS predictions over the number of predicted CS, recall as the fraction of correct CS predictions over the number of true CS. In both cases, a CS was only considered correct if it was predicted in the correct SP class (e.g. when the model predicts a CS in a Sec/SPI sequence, but predicts Sec/SPII as the sequence label, the sample is considered as no CS predicted). To account for possible uncertainty of the CS in the training data labels, we additionally report these metrics with tolerance windows of 1, 2 and 3 residues left and right of the true CS (Tables S4, S6, S8).

For the predicted SP regions, in the absence of true labels, no quantitative performance metrics could be established. To still be able to assess the quality of the predictions, we compared the properties of predicted regions to characteristics of regions that are described in literature. We followed the review by Owji et al. (18) as a guideline to identify region characteristics. Specifically, we evaluated the length, hydrophobicity, and charge of each predicted region. Hydrophobicities were computed using the Kyte-Doolittle scale (31), charges by summing the net charges at pH 7 of all residues. The net charge computation differed between the groups, as in Eukarya and Archaea the N-terminal methionine is not formylated (35), thus contributing an additional positive charge to the n-region by its amino group.

We benchmarked our model against the state-of-the-art model SignalP 5.0, which was reimplemented in Pytorch. Hyperparameter optimization on the new dataset was performed using SigOpt. We also repeated the benchmarking experiment of SignalP 5.0 for all predictors using the adapted benchmark set. We could not add Signal-3L 3.0 (16) to the experiment, as the implementation that is available does not allow for processing of more than one sequence at a time, rendering benchmarking intractable. Notably, predictions for all methods except for SignalP 5.0 and SignalP 6.0 were obtained from their publicly available web services, resulting in potential performance overestimation due to the lack of homology partitioning. Additionally, performance overestimation is still present for the published version of SignalP 5.0 (named “SignalP 5.0 original” in Tables S3-S9) due to insufficient homology partitioning of its training data by CD-HIT. We thus excluded its values from determining the best performing tools in the benchmark.

To assess the effect of sequence identity to training sequences on performance, we used the set of sequences that were removed by the partitioning procedure. We predicted all sequences in the removed set and binned the sequences according to the maximum sequence identity to any sequence in the training set. We did this for all 6 cross-validated models and pooled the resulting binned predictions. For each bin, we computed the multi-class MCC as defined by Gorodkin (36).

