## Supplementary figures S1-S4, Supplementary tables S1-S11, Supplementary Text for "SignalP 6.0 achieves signal peptide prediction across all types using protein language models"

Model distillation

The SignalP 6.0 predictor is an ensemble model of all six models trained in nested cross-validation. Due to the computational demand of the BERT protein LM, this poses a challenge for prediction on systems with limited resources and results in long processing times for larger numbers of sequences. To still be able to offer prediction speeds comparable to previous versions of SignalP, we performed model distillation. For each sequence in the training set, we obtain the marginal probabilities predicted by the full ensemble model (termed *SignalP 6.0 - slow*). We instantiate a new SignalP 6.0 model from the pretrained LM, and train this model on the full training set, using the Kullback-Leibler divergence between its marginal probabilities and the ensemble model marginal probabilities as the loss. To mitigate class imbalance, we multiply the loss of each sequence with a weight inversely proportional to its type's prevalence in the training set. The model is trained until convergence, yielding *SignalP 6.0 - fast*.

By nature of the training objective, the distilled model is only trained to approximate the marginal probabilities and SP type probabilities of the full model (Table S11). However, as the distilled model does not replicate the emissions, this can lead to discrepancies in the predicted Viterbi paths. We thus recommend using *SignalP 6.0 - slow* for applications where accurate region border predictions are of interest.

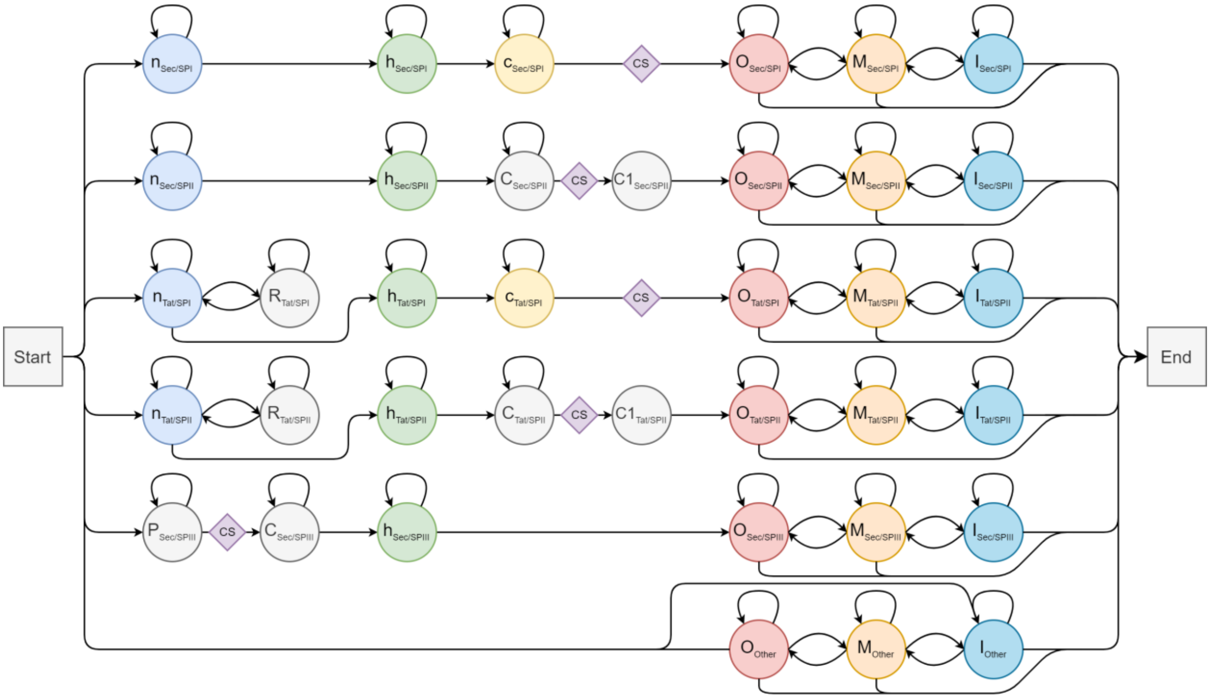


**Figure S1.** States modeled by the CRF. The three regions are indicated by their lower-case name. R is the state of the twin-arginine motif. In lipoproteins, C marks the lipobox and C1 marks the cysteine in +1 of the cleavage site. CS indicates the position of the cleavage site, which is not modeled as a state, but inferred from the end of the previous region. For Sec/SPIII, the whole SP is modeled as a single state P, followed by a conserved and a hydrophobic region. O, M, and I mark extracellular, transmembrane and intracellular regions of the mature protein.

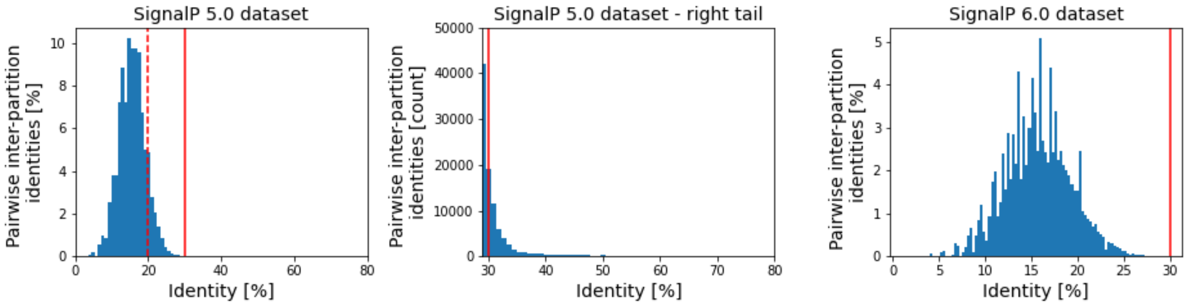


**Figure S2.** Quality of homology partitioning. All pairwise inter-partition identities were computed using ggsearch36. The dashed line indicates the reported threshold for SignalP 5.0, the solid line the relaxed threshold of 30% used in this work.

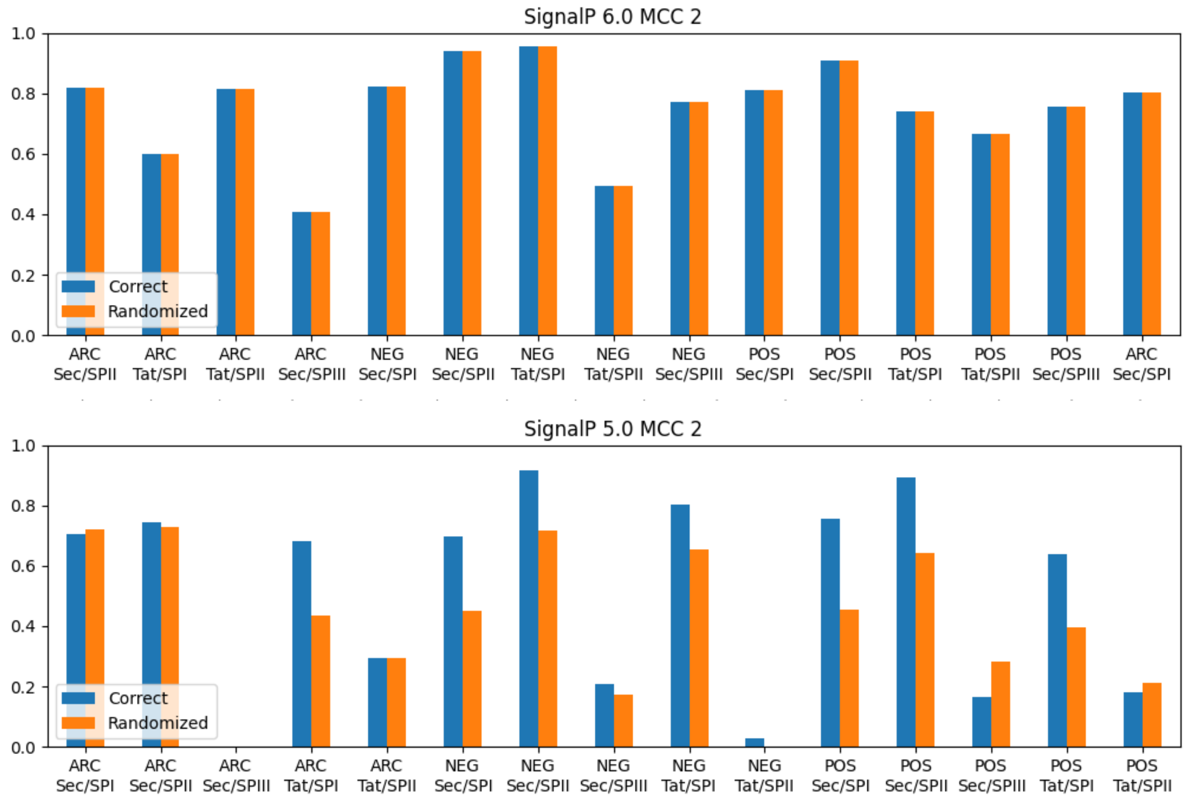


**Figure S3.** Performance of SignalP 5.0 and SignalP 6.0 on data with correct and with randomized organism group identifiers.

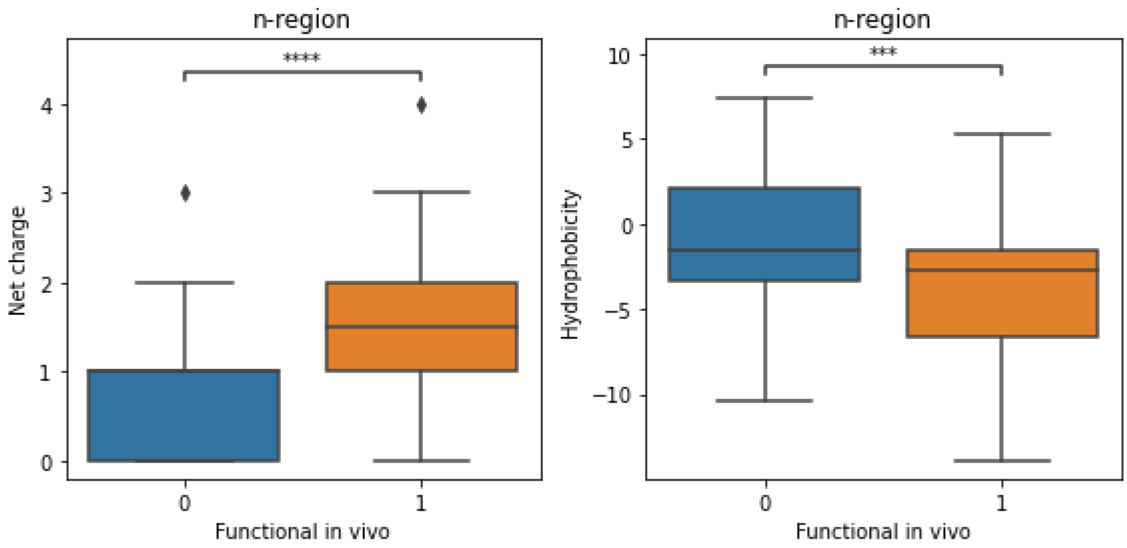


**Figure S4.** Comparison of n-region properties of synthetic SPs. In *B. subtilis*, group 0 was found to be nonfunctional, for group 1 protein secretion was observed (****: $p<1\times10^{-4}$, ***: $p<1\times10^{-3}$, Welch's t-test).

| | Eukarya | Archaea | Gram-positive | Gram-negative |
|---|---|---|---|---|
| Sec/SPI | 2040 (2840) | 44 (61) | 142 (214) | 356 (537) |
| Sec/SPII | - | 12 (27) | 516 (782) | 1087 (1452) |
| Sec/SPIII | - | 10 (10) | 4 (6) | 56 (97) |
| Tat/SPI | - | 13 (24) | 39 (110) | 313 (461) |
| Tat/SPII | - | 6 (6) | 8 (11) | 19 (19) |
| Other | 14356 (17627) | 110 (124) | 226 (258) | 933 (1077) |

**Table S1.** Composition of the SignalP 6.0 training set. Numbers in parentheses are counts before application of the homology partitioning procedure.

| | Eukarya | Archaea | Gram-negative bacteria | Gram-positive bacteria |
|---|---|---|---|---|
| Sec/SPI | 146 | 36 | 61 | 15 |
| Sec/SPII | - | 9 | 257 | 120 |
| Tat/SPI | - | 9 | 51 | 18 |
| Tat/SPII | - | 5 | 5 | 3 |
| Other | 5581 | 81 | 133 | 81 |

**Table S2.** Composition of the SignalP 5.0 benchmark dataset after 1) removal of all sequences that were not retained in the new homology partitioning and 2) reclassification of Gram-negative and Tat/SPI samples to Gram-positive and Tat/SPII.

| Method | Archaea | | Eukarya | Gram-negative bacteria | | Gram-positive bacteria | |
|---|---|---|---|---|---|---|---|
| | MCC1 | MCC2 | MCC1 | MCC1 | MCC2 | MCC1 | MCC2 |
| SignalP 6.0 | 0.737 | **0.728** | **0.868** | 0.811 | **0.649** | 0.878 | **0.734** |
| SignalP 5.0 retrained | 0.711 | 0.67 | 0.774 | 0.705 | 0.586 | 0.798 | 0.669 |
| DEEPSIG | n.d. | n.d. | 0.792 | 0.735 | 0.159 | 0.798 | 0.146 |
| LipoP | 0.775 | 0.619 | 0.347 | 0.744 | 0.471 | **0.879** | 0.442 |
| PHILIUS | 0.691 | 0.438 | 0.448 | 0.766 | 0.147 | 0.752 | 0.084 |
| PHOBIUS | 0.796 | 0.551 | 0.531 | 0.766 | 0.153 | 0.716 | 0.08 |
| PolyPhobius | 0.715 | 0.474 | 0.478 | 0.813 | 0.173 | 0.777 | 0.136 |
| PRED-LIPO | 0.733 | 0.552 | 0.196 | 0.71 | 0.342 | 0.879 | 0.484 |
| PRED-SIGNAL | **0.908** | 0.67 | 0.265 | 0.662 | 0.115 | 0.822 | 0.171 |
| PRED-TAT | 0.781 | 0.655 | 0.34 | 0.736 | 0.209 | 0.839 | 0.238 |
| SIGNAL-CF | n.d. | n.d. | 0.333 | 0.52 | 0.123 | 0.474 | 0.1 |
| Signal-3L 2.0 | n.d. | n.d. | 0.605 | 0.731 | 0.108 | 0.878 | 0.133 |
| SOSUIsignal | n.d. | n.d. | 0.368 | 0.639 | 0.123 | 0.702 | 0.107 |
| SPEPlip | n.d. | n.d. | 0.652 | 0.705 | 0.489 | 0.578 | 0.429 |
| SPOCTOPUS | 0.732 | 0.448 | 0.506 | **0.849** | 0.165 | **0.879** | 0.134 |
| TOPCONS2 | 0.711 | 0.438 | 0.504 | 0.844 | 0.159 | 0.836 | 0.078 |
| SignalP 5.0 original | 0.899 | 0.886 | 0.863 | 0.821 | 0.77 | 0.921 | 0.868 |

**Table S3.** Benchmark results for Sec/SPI prediction. MCC1 refers to detection performance when the negative class consists of soluble and transmembrane proteins. For MCC2, the negative class additionally contains Sec/SPII, Tat/SPI and Tat/SPII SPs.

| Method | Archaea | | | | Eukarya | | | | Gram-negative bacteria | | | | Gram-positive bacteria | | | |
|---|---|---|---|---|---|---|---|---|---|---|---|---|---|---|---|---|
| | ±0 | ±1 | ±2 | ±3 | ±0 | ±1 | ±2 | ±3 | ±0 | ±1 | ±2 | ±3 | ±0 | ±1 | ±2 | ±3 |
| CS recall | | | | | | | | | | | | | | | | |
| SignalP 6.0 | 0.500 | 0.556 | 0.556 | 0.583 | **0.747** | **0.774** | **0.808** | **0.829** | 0.639 | 0.672 | 0.689 | 0.721 | 0.800 | 0.800 | 0.800 | 0.800 |
| SignalP 5.0 retrained | 0.389 | 0.472 | 0.472 | 0.528 | 0.63 | 0.651 | 0.705 | 0.760 | 0.508 | 0.574 | 0.656 | 0.672 | 0.733 | 0.733 | 0.733 | 0.733 |
| DEEPSIG | n.d. | n.d. | n.d. | n.d. | 0.603 | 0.63 | 0.658 | 0.699 | 0.508 | 0.574 | 0.574 | 0.574 | 0.733 | 0.733 | 0.800 | 0.800 |
| LipoP | 0.389 | 0.528 | 0.556 | 0.639 | 0.288 | 0.329 | 0.370 | 0.404 | 0.656 | 0.705 | 0.721 | 0.721 | 0.467 | 0.467 | 0.533 | 0.533 |
| PHILIUS | 0.500 | 0.611 | 0.611 | 0.611 | 0.596 | 0.658 | 0.712 | 0.760 | 0.623 | 0.672 | 0.721 | 0.754 | 0.467 | 0.467 | 0.467 | 0.467 |
| PHOBIUS | 0.472 | 0.583 | 0.611 | 0.639 | 0.637 | 0.671 | 0.699 | 0.753 | 0.557 | 0.656 | 0.721 | 0.738 | 0.467 | 0.467 | 0.467 | 0.467 |
| PolyPhobius | 0.528 | 0.667 | 0.667 | 0.667 | 0.623 | 0.678 | 0.733 | 0.801 | 0.557 | 0.672 | 0.754 | 0.754 | 0.667 | 0.667 | 0.733 | 0.733 |
| PRED-LIPO | 0.472 | 0.556 | 0.611 | 0.639 | 0.068 | 0.082 | 0.130 | 0.158 | 0.410 | 0.475 | 0.508 | 0.525 | **0.867** | **0.867** | **0.867** | **0.867** |
| PRED-SIGNAL | **0.861** | **0.917** | **0.917** | **0.917** | 0.226 | 0.267 | 0.301 | 0.329 | 0.426 | 0.492 | 0.607 | 0.639 | 0.800 | 0.800 | 0.800 | 0.800 |
| PRED-TAT | 0.556 | 0.694 | 0.75 | 0.778 | 0.370 | 0.445 | 0.500 | 0.548 | **0.656** | **0.721** | 0.754 | 0.770 | **0.867** | **0.867** | **0.867** | **0.867** |
| Signal-3L 2.0 | n.d. | n.d. | n.d. | n.d. | 0.644 | 0.671 | 0.719 | 0.753 | 0.607 | 0.639 | 0.672 | 0.705 | 0.733 | 0.733 | 0.800 | 0.800 |
| Signal3Lv2 | n.d. | n.d. | n.d. | n.d. | 0.664 | 0.685 | 0.726 | 0.753 | 0.541 | 0.607 | 0.623 | 0.639 | 0.800 | 0.800 | 0.800 | 0.800 |
| SOSUIsignal | n.d. | n.d. | n.d. | n.d. | 0.151 | 0.308 | 0.459 | 0.568 | 0.246 | 0.377 | 0.557 | 0.623 | 0.200 | 0.267 | 0.267 | 0.467 |
| SPEPlip | n.d. | n.d. | n.d. | n.d. | 0.685 | 0.712 | 0.747 | 0.781 | 0.574 | 0.656 | 0.705 | 0.721 | 0.600 | 0.600 | 0.667 | **0.667** |
| SPOCTOPUS | 0.333 | 0.389 | 0.417 | 0.472 | 0.384 | 0.514 | 0.678 | 0.747 | 0.426 | 0.656 | **0.820** | **0.869** | 0.600 | 0.667 | 0.733 | **0.867** |
| TOPCONS2 | 0.389 | 0.528 | 0.556 | 0.583 | 0.329 | 0.452 | 0.596 | 0.692 | 0.443 | 0.541 | 0.656 | 0.689 | 0.267 | 0.333 | 0.333 | 0.400 |
| SignalP 5.0 original | 0.611 | 0.694 | 0.722 | 0.778 | 0.692 | 0.740 | 0.767 | 0.815 | 0.672 | 0.705 | 0.738 | 0.738 | 0.933 | 0.933 | 0.933 | 0.933 |
| CS precision | | | | | | | | | | | | | | | | |
| SignalP 6.0 | **0.643** | **0.714** | **0.714** | **0.75** | **0.661** | **0.685** | **0.715** | **0.733** | **0.534** | **0.562** | **0.575** | **0.603** | **0.632** | **0.632** | **0.632** | **0.632** |
| SignalP 5.0 retrained | 0.519 | 0.630 | 0.630 | 0.704 | 0.514 | 0.531 | 0.575 | 0.620 | 0.378 | 0.427 | 0.488 | 0.500 | 0.500 | 0.500 | 0.500 | 0.500 |
| DEEPSIG | n.d. | n.d. | n.d. | n.d. | 0.587 | 0.613 | 0.640 | 0.680 | 0.134 | 0.151 | 0.151 | 0.151 | 0.089 | 0.089 | 0.098 | 0.098 |
| LipoP | 0.359 | 0.487 | 0.513 | 0.590 | 0.141 | 0.162 | 0.182 | 0.199 | 0.339 | 0.364 | 0.373 | 0.373 | 0.152 | 0.152 | 0.174 | 0.174 |
| PHILIUS | 0.353 | 0.431 | 0.431 | 0.431 | 0.168 | 0.186 | 0.201 | 0.215 | 0.110 | 0.118 | 0.127 | 0.133 | 0.051 | 0.051 | 0.051 | 0.051 |
| PHOBIUS | 0.340 | 0.420 | 0.440 | 0.460 | 0.245 | 0.258 | 0.268 | 0.289 | 0.099 | 0.117 | 0.129 | 0.132 | 0.051 | 0.051 | 0.051 | 0.051 |
| PolyPhobius | 0.352 | 0.444 | 0.444 | 0.444 | 0.181 | 0.197 | 0.213 | 0.233 | 0.098 | 0.118 | 0.133 | 0.133 | 0.069 | 0.069 | 0.076 | 0.076 |
| PRED-LIPO | 0.386 | 0.455 | 0.500 | 0.523 | 0.052 | 0.062 | 0.098 | 0.119 | 0.203 | 0.236 | 0.252 | 0.26 | 0.325 | 0.325 | 0.325 | 0.325 |
| PRED-SIGNAL | 0.508 | 0.541 | 0.541 | 0.541 | 0.073 | 0.086 | 0.097 | 0.106 | 0.085 | 0.098 | 0.121 | 0.128 | 0.083 | 0.083 | 0.083 | 0.083 |
| PRED-TAT | 0.426 | 0.532 | 0.574 | 0.596 | 0.08 | 0.097 | 0.109 | 0.119 | 0.133 | 0.147 | 0.153 | 0.157 | 0.101 | 0.101 | 0.101 | 0.101 |
| Signal-3L 2.0 | n.d. | n.d. | n.d. | n.d. | 0.103 | 0.108 | 0.115 | 0.121 | 0.104 | 0.110 | 0.115 | 0.121 | 0.067 | 0.067 | 0.074 | 0.074 |
| Signal3Lv2 | n.d. | n.d. | n.d. | n.d. | 0.357 | 0.368 | 0.39 | 0.404 | 0.116 | 0.130 | 0.134 | 0.137 | 0.093 | 0.093 | 0.093 | 0.093 |
| SOSUIsignal | n.d. | n.d. | n.d. | n.d. | 0.032 | 0.066 | 0.098 | 0.122 | 0.042 | 0.065 | 0.096 | 0.107 | 0.021 | 0.028 | 0.028 | 0.049 |
| SPEPlip | n.d. | n.d. | n.d. | n.d. | 0.362 | 0.377 | 0.395 | 0.413 | 0.278 | 0.317 | 0.341 | 0.349 | 0.257 | 0.257 | 0.286 | 0.286 |
| SPOCTOPUS | 0.240 | 0.280 | 0.300 | 0.340 | 0.127 | 0.170 | 0.224 | 0.247 | 0.070 | 0.107 | 0.134 | 0.142 | 0.062 | 0.068 | 0.075 | 0.089 |
| TOPCONS2 | 0.275 | 0.373 | 0.392 | 0.412 | 0.110 | 0.151 | 0.199 | 0.231 | 0.078 | 0.095 | 0.115 | 0.121 | 0.029 | 0.036 | 0.036 | 0.043 |
| SignalP 5.0 original | 0.647 | 0.735 | 0.765 | 0.824 | 0.635 | 0.679 | 0.704 | 0.748 | 0.719 | 0.754 | 0.789 | 0.789 | 0.824 | 0.824 | 0.824 | 0.824 |

**Table S4.** Benchmark results for CS prediction in Sec/SPI at different tolerance windows

| Method | Archaea | | Gram-negative bacteria | | Gram-positive bacteria | |
|---|---|---|---|---|---|---|
| | MCC1 | MCC2 | MCC1 | MCC2 | MCC1 | MCC2 |
| SignalP 6.0 | **0.871** | **0.719** | 0.838 | 0.841 | **0.894** | **0.893** |
| SignalP 5.0 retrained | **0.871** | **0.719** | **0.884** | **0.874** | 0.883 | 0.866 |
| LipoP | **0.871** | 0.681 | 0.806 | 0.813 | 0.71 | 0.724 |
| PRED-LIPO | 0.728 | 0.608 | 0.615 | 0.655 | 0.762 | 0.743 |
| SPEPlip | n.d. | n.d. | 0.856 | 0.86 | 0.842 | 0.837 |
| SignalP 5.0 original | 0.937 | 0.881 | 0.939 | 0.925 | 0.922 | 0.917 |

**Table S5.** Benchmark results for Sec/SPII prediction. MCC1 refers to detection performance when the negative class consists of soluble and transmembrane proteins. For MCC2, the negative class additionally contains Sec/SPI, Tat/SPI and Tat/SPII SPs.

| Method | Archaea | | | | Gram-negative bacteria | | | | Gram-positive bacteria | | | |
|---|---|---|---|---|---|---|---|---|---|---|---|---|
| | ±0 | ±1 | ±2 | ±3 | ±0 | ±1 | ±2 | ±3 | ±0 | ±1 | ±2 | ±3 |
| **CS recall** | | | | | | | | | | | | |
| SignalP 6.0 | **0.778** | **0.778** | **0.778** | **0.778** | 0.852 | 0.852 | 0.856 | 0.864 | 0.875 | 0.883 | 0.883 | 0.883 |
| SignalP 5.0 retrained | **0.778** | **0.778** | **0.778** | **0.778** | **0.895** | **0.895** | **0.895** | **0.907** | **0.900** | **0.900** | **0.900** | **0.900** |
| LipoP | **0.778** | **0.778** | **0.778** | **0.778** | 0.837 | 0.837 | 0.837 | 0.837 | 0.700 | 0.700 | 0.700 | 0.700 |
| PRED-LIPO | 0.556 | 0.556 | 0.556 | 0.556 | 0.646 | 0.646 | 0.646 | 0.646 | 0.767 | 0.767 | 0.767 | 0.767 |
| SPEPlip | n.d. | n.d. | n.d. | n.d. | 0.887 | 0.887 | 0.891 | 0.891 | 0.850 | 0.850 | 0.850 | 0.850 |
| SignalP 5.0 original | 0.889 | 0.889 | 0.889 | 0.889 | 0.949 | 0.949 | 0.949 | 0.953 | 0.917 | 0.917 | 0.917 | 0.917 |
| **CS precision** | | | | | | | | | | | | |
| SignalP 6.0 | 0.583 | 0.583 | 0.583 | 0.583 | 0.913 | 0.913 | 0.917 | 0.925 | 0.929 | 0.938 | 0.938 | 0.938 |
| SignalP 5.0 retrained | 0.583 | 0.583 | 0.583 | 0.583 | 0.895 | 0.895 | 0.895 | 0.907 | 0.931 | 0.931 | 0.931 | 0.931 |
| LipoP | 0.636 | 0.636 | 0.636 | 0.636 | 0.951 | 0.951 | 0.951 | 0.951 | 0.955 | 0.955 | 0.955 | 0.955 |
| PRED-LIPO | **0.714** | **0.714** | **0.714** | **0.714** | **0.954** | **0.954** | 0.954 | 0.954 | 0.939 | 0.939 | 0.939 | 0.939 |
| SPEPlip | n.d. | n.d. | n.d. | n.d. | 0.954 | 0.954 | **0.958** | **0.958** | **0.962** | **0.962** | **0.962** | **0.962** |
| SignalP 5.0 original | 0.889 | 0.889 | 0.889 | 0.889 | 0.953 | 0.953 | 0.953 | 0.957 | 0.965 | 0.965 | 0.965 | 0.965 |

**Table S6.** Benchmark results for CS prediction in Sec/SPII at different tolerance windows.

| Method | Archaea | | Gram-negative bacteria | | Gram-positive bacteria | |
|---|---|---|---|---|---|---|
| | MCC1 | MCC2 | MCC1 | MCC2 | MCC1 | MCC2 |
| SignalP 6.0 | 0.802 | **0.807** | **0.946** | **0.934** | 0.788 | **0.806** |
| SignalP 5.0 retrained | 0.807 | 0.763 | 0.719 | 0.732 | 0.708 | 0.700 |
| PRED-TAT | 0.937 | 0.719 | 0.945 | 0.869 | **0.823** | 0.643 |
| TatP | 0.733 | 0.474 | 0.730 | 0.591 | 0.568 | 0.411 |
| TATFIND | **0.937** | 0.662 | 0.892 | 0.845 | 0.711 | 0.580 |
| SignalP 5.0 original | 0.937 | 0.719 | 0.973 | 0.934 | 0.931 | 0.880 |

**Table S7.** Benchmark results for Tat/SPI prediction. MCC1 refers to detection performance when the negative class consists of soluble and transmembrane proteins. For MCC2, the negative class additionally contains Sec/SPI, Sec/SPII and Tat/SPII SPs.

| Method | Archaea | | | | Gram-negative bacteria | | | | Gram-positive bacteria | | | |
|---|---|---|---|---|---|---|---|---|---|---|---|---|
| | ±0 | ±1 | ±2 | ±3 | ±0 | ±1 | ±2 | ±3 | ±0 | ±1 | ±2 | ±3 |
| **CS recall** | | | | | | | | | | | | |
| SignalP 6.0 | **0.333** | **0.444** | 0.444 | 0.444 | 0.706 | **0.765** | **0.784** | 0.804 | 0.556 | 0.556 | **0.667** | 0.667 |
| SignalP 5.0 retrained | 0.222 | 0.444 | 0.444 | 0.444 | 0.412 | 0.451 | 0.490 | 0.490 | 0.167 | 0.222 | 0.222 | 0.278 |
| PRED-TAT | **0.333** | **0.444** | **0.667** | **0.667** | **0.725** | **0.765** | **0.784** | **0.824** | **0.611** | **0.611** | **0.667** | **0.722** |
| TatP | 0.222 | 0.333 | 0.444 | 0.444 | 0.588 | 0.608 | 0.608 | 0.627 | 0.333 | 0.333 | 0.389 | 0.389 |
| TATFIND | n.d. | n.d. | n.d. | n.d. | n.d. | n.d. | n.d. | n.d. | n.d. | n.d. | n.d. | n.d. |
| SignalP 5.0 original | 0.333 | 0.444 | 0.556 | 0.556 | 0.686 | 0.745 | 0.784 | 0.804 | 0.667 | 0.667 | 0.833 | 0.833 |
| **CS precision** | | | | | | | | | | | | |
| SignalP 6.0 | **0.375** | **0.500** | **0.500** | **0.500** | **0.679** | **0.736** | **0.755** | **0.774** | **0.714** | **0.714** | **0.857** | **0.857** |
| SignalP 5.0 retrained | 0.182 | 0.364 | 0.364 | 0.364 | 0.488 | 0.535 | 0.581 | 0.581 | 0.273 | 0.364 | 0.364 | 0.455 |
| PRED-TAT | 0.231 | 0.308 | 0.462 | 0.462 | 0.638 | 0.672 | 0.690 | 0.724 | 0.458 | 0.458 | 0.500 | 0.542 |
| TatP | 0.133 | 0.200 | 0.267 | 0.267 | 0.326 | 0.337 | 0.337 | 0.348 | 0.167 | 0.167 | 0.194 | 0.194 |
| TATFIND | n.d. | n.d. | n.d. | n.d. | n.d. | n.d. | n.d. | n.d. | n.d. | n.d. | n.d. | n.d. |
| SignalP 5.0 original | 0.231 | 0.308 | 0.385 | 0.385 | 0.660 | 0.717 | 0.755 | 0.774 | 0.667 | 0.667 | 0.833 | 0.833 |

**Table S8.** Benchmark results for CS prediction in Tat/SPI at different tolerance windows

| Species name | Taxonomy ID | Other | Sec/SPI | Sec/SPII | Sec/SPIII | Tat/SPI | Tat/SPII | Note |
|---|---|---|---|---|---|---|---|---|
| Methanosaeta harundinacea | 1110509 | 2065 | 260 | 33 | 0 | 0 | 0 | Max. Sec/SPI frequency in Archaea |
| Methanolacinia petrolearia | 679926 | 2511 | 163 | 97 | 8 | 0 | 0 | Max. Sec/SPII frequency in Archaea |
| Halorussus sp. MSC15.2 | 2283638 | 3704 | 94 | 24 | 2 | 141 | 107 | Max. Tat/SPI frequency in Archaea |
| Natrialba swarupiae | 2448032 | 3585 | 43 | 11 | 2 | 43 | 133 | Max. Tat/SPII frequency in Archaea |
| Candidate division MSBL1 archaeon SCGC-AAA382A13 | 1698279 | 439 | 6 | 0 | 5 | 0 | 0 | Max. Sec/SPIII frequency in Archaea |
| Acidilobus sp. SCGC AC-742_M05 | 1987489 | 280 | 1 | 0 | 0 | 0 | 0 | Min. Sec/SPI frequency in Archaea |
| archaeon HR04 | 2035440 | 1311 | 50 | 0 | 0 | 1 | 0 | Min. Sec/SPII frequency in Archaea |
| Methanoplanus limicola DSM 2279 | 937775 | 2666 | 166 | 88 | 8 | 0 | 0 | Min. Tat/SPI frequency in Archaea |
| Candidate division TM6 bacterium JCVI TM6SC1 | 1306947 | 526 | 304 | 24 | 6 | 0 | 0 | Max. Sec/SPI frequency in Bacteria |
| Nannocystis exedens | 54 | 6725 | 669 | 1664 | 5 | 55 | 25 | Max. Sec/SPII frequency in Bacteria |
| Roseomonas stagni DSM 19981 | 1123062 | 4961 | 403 | 109 | 0 | 421 | 8 | Max. Tat/SPI frequency in Bacteria |
| Eggerthella sp. (strain YY7918) | 502558 | 2476 | 51 | 44 | 5 | 14 | 84 | Max. Tat/SPII frequency in Bacteria |
| Victivallis vadensis | 172901 | 3167 | 493 | 184 | 210 | 5 | 0 | Max. Sec/SPIII frequency in Bacteria |
| Buchnera aphidicola (Stegophylla sp.) | 2315800 | 354 | 0 | 0 | 0 | 0 | 0 | Bacterial endosymbiont |
| Mycoplasma canadense | 29554 | 426 | 0 | 55 | 0 | 0 | 0 | Min. Sec/SPI frequency in Bacteria |
| Candidatus Mikella endobia | 1778264 | 272 | 1 | 0 | 0 | 0 | 0 | Bacterial endosymbiont,<br>Min. Sec/SPII frequency in Bacteria |
| Candidatus Termititenax dinenymphae | 2218523 | 335 | 10 | 10 | 0 | 0 | 0 | Min. Tat/SPI frequency in Bacteria |
| Escherichia coli (strain K12) | 83333 | 3854 | 378 | 123 | 8 | 27 | 1 | |
| Thermus thermophilus | 300852 | 2028 | 132 | 39 | 9 | 18 | 1 | |
| Deinococcus radiodurans | 243230 | 2744 | 238 | 91 | 4 | 7 | 1 | |
| Corynebacterium glutamicum | 196627 | 2887 | 99 | 94 | 0 | 3 | 10 | |
| Thermotoga maritima | 243274 | 1706 | 121 | 21 | 4 | 0 | 0 | |
| Bacillus subtilis | 224308 | 3988 | 151 | 113 | 4 | 4 | 0 | |
| Archaeoglobus veneficus | 693661 | 1954 | 59 | 41 | 4 | 5 | 2 | |
| Haloferax volcanii | 309800 | 3699 | 57 | 16 | 3 | 31 | 105 | |
| Methanocaldococcus jannaschii | 243232 | 1717 | 31 | 28 | 11 | 0 | 0 | |
| Prometheoarchaeum syntrophicum | 2594042 | 3813 | 102 | 18 | 0 | 0 | 0 | |
| Pyrococcus furiosus | 186497 | 1937 | 75 | 29 | 4 | 0 | 0 | |

**Table S9.** Selected UniProt reference proteome predictions. Species with no SPs predicted (n=9) were excluded for the determination of bacteria with minimum frequencies.

| Organism group | % Sec/SPI | % Sec/SPII | % Sec/SPIII | % Tat/SPI | % Tat/SPII |
|---|---|---|---|---|---|
| Archaea (n=330) | 2.65 ± 1.96 | 0.58 ± 0.66 | 0.12 ± 0.17 | 0.43 ± 0.62 | 0.71 ± 1.05 |
| Eukarya (n=1588) | 8.08 ± 3.97 | - | - | - | - |
| Gram-positive (n=3387) | 3.10 ± 1.43 | 2.78 ± 1.2 | 0.11 ± 0.16 | 0.21 ± 0.29 | 0.14 ± 0.27 |
| Gram-negative (n=4610) | 9.84 ± 3.87 | 3.98 ± 2.87 | 0.24 ± 0.38 | 0.73 ± 0.71 | 0.24 ± 0.10 |

**Table S10.** Average frequencies and standard deviations of the five signal peptide types in UniProt reference proteomes.

| True \ Pred. | Other | Sec/SPI | Sec/SPII | Sec/SPIII | Tat/SPI | Tat/SPII |
|---|---|---|---|---|---|---|
| **Other** | 3393 (3392) | 14 (16) | 2 (0) | 1 (1) | 1 (2) | 0 (0) |
| **Sec/SPI** | 12 (8) | 1046 (1056) | 11 (5) | 0 (0) | 1 (1) | 0 (0) |
| **Sec/SPII** | 1 (1) | 6 (7) | 638 (637) | 0 (0) | 0 (0) | 1 (1) |
| **Sec/SPIII** | 0 (0) | 0 (0) | 0 (0) | 43 (43) | 0 (0) | 0 (0) |
| **Tat/SPI** | 1 (0) | 10 (15) | 1 (0) | 0 (0) | 213 (206) | 5 (9) |
| **Tat/SPII** | 0 (0) | 0 (0) | 0 (0) | 0 (0) | 0 (0) | 3 (3) |

| Class | MCC 2 |
|---|---|
| Other | 0.982 (0.989) |
| Sec/SPI | 0.987 (0.989) |
| Sec/SPII | 0.988 (0.988) |
| Sec/SPIII | 0.969 (0.970) |
| Tat/SPI | 0.956 (0.957) |
| Tat/SPII | 0.577 (0.480) |

**Table S11.** Comparison of the distilled model to the full ensemble model, using the set of sequences that were removed by the partitioning procedure as the test set. The performance of the full ensemble model is given in parentheses.
